# Cell phone digital microscopy using an oil droplet

**DOI:** 10.1101/2020.02.10.942490

**Authors:** Nicole Anna Szydlowski, Haoran Jing, Mohamed Alqashmi, Ying Samuel Hu

## Abstract

We introduce an accessible imaging method using droplets of microscope immersion and consumer-grade oils and a cell phone camera. We found that oil droplets were more resistant to evaporation than water droplets. We characterized the transverse magnification of oil and water droplets using cell phone screens and a resolution target. We further harnessed the close refractive index of cooking oils to that of the immersion oil and demonstrated their use as lenses for cell phone microscopy. Our method enables stable droplet-based optical imaging without specialized setups or manufacturing processes.

## 1. Introduction

Cutting-edge optical microscopy is currently in high demand in the fields of medicine and biology research. Nevertheless, in low-resource settings where accessibility is limited, the ability to quickly assess the morphology and size of the biological specimen beyond what the human eye can see is of practical interest. In response to the demand for access to low-cost microscopy for educational and diagnostic purposes, several researchers have developed new microscopy devices by attaching lenses and other types of devices to smart phones to perform brightfield, darkfield, fluorescence, and polarized imaging ^1–7^. An added benefit of smart phones is that they can be used to transmit high-quality images through Multimedia Messaging Services (MMS). As of 2007-2008, the percentage of people who had MMS in countries across Africa ranged from 1.5% to 92.2% ^8^. This increasingly widespread cellular connectivity could be harnessed to facilitate more rapid scientific communication between individuals and to increase accessibility to microscopy for diagnostic and educational purposes on demand.

Previous studies have demonstrated the ability to capture clinically useful microscopic images using a ball lens in front of the camera lens of a cell phone ^9,10^. One study developed on-demand lenses by heat-curing polydimethylsiloxane (PDMS) plano-convex lenses in order to conduct cell phone imaging without using attachments such as ball lenses or accessory devices ^11^. While uncured PDMS becomes too thin to function as an effective lens, heat-cured PDMS maintains its droplet shape well, which has been shown to enable a magnification of up to 120 times and a resolution of up to 1 micron ^11^. Other studies have explored the option of creating tunable liquid lenses whose focal lengths are tunable through variations in pressure distribution in a liquid-containing chamber using a temperature-sensitive or pH-sensitive hydrogel ring ^12^. While these inexpensive lenses enable a vast range of imaging applications with ease of operation, the distribution channels of custom-made lenses to low-resource areas have become a major bottleneck. Moreover, increasing numbers of custom applications call for the on-demand design of accessible, cost-effective lenses. Here, we seek to evaluate the use of simple and accessible materials for optical imaging.

Water droplets are easy to make and do not require specialized fabrication processes, so they can serve as useful tools for microscopy ^13^. Nevertheless, water droplets have two significant limitations for imaging applications. First, water droplets evaporate rapidly under ambient conditions, which changes their focal length over time. To achieve optical amplification, water droplets often have small volumes, *i.e.*, less than 10 µl. Temperature, air flow, and humidity affect the evaporation rate, which quickly diminishes the optical magnification. Some studies have used methods to reduce the rate of water droplet evaporation, which enables longer imaging sessions while maintaining a consistent focal length. However, even with these methods, water droplet imaging methods are still very time-limited due to water evaporation. One study used a plastic container with wet paper next to the water droplet to maintain a consistent water vapor pressure, which maintained a constant focal length in the water droplet for two hours ^13^. Another study used spherical water droplets at the tip of a syringe needle as lenses and coated them with a silicone oil to reduce evaporation so that the water droplets could be used for an hour ^14^. While each of these studies succeeded in developing a more flexible approach to water droplet microscopy, it may be difficult to conduct certain microscopy experiments using only a one to two hour working time. Therefore, one of our focuses was to develop a method that would enable liquid droplets to be used over a much longer period of time. In addition to evaporating quickly, water-based lenses display optical aberration due to the index mismatch between water and glass. Since most biological specimens are mounted on a coverglass, which has a refractive index of 1.515 compared to 1.33 of water, optical refraction at the interface could deteriorate the image quality. In this report, we investigate the use of oil droplets that can be used in smartphone microscopy to obtain images of biological samples.

We started by demonstrating the use of index-matched immersion oil droplets for stable optical imaging, and then extend the method by using household cooking oils. We obtained the refractive index values for common household liquids from the International Gem Society ^15^. For instance, safflower, peanut, and sesame oil have refractive indices around 1.47-1.48, closely resembling the refractive index of immersion oil at 1.515 ^16^. Palm oil has a slightly lower refractive index of 1.46-1.47 ^17^. In this study, we decided to compare droplets using corn oil, canola oil, and olive oil.

## 2. Methods

### 2.1 Droplet magnification analysis

To characterize the optical amplification of droplets, we first used cell phone screens for optical illumination and measured amplified pixels through the droplet on a cellphone screen (Fig 1). To prepare a series of droplet “lenses,” we used a micropipette to place droplets with volumes ranging from 1-5 µL in a row on a No. 1.5 (170±5 µm) coverglass. For precise pipetting of the oil droplet, we prewarmed the immersion oil (Nikon Immersion Oil Type F, index = 1.518) in a 37°C water bath to reduce its viscosity. We lifted the coverglass above the cell phone screen through a stack of three glass slides measuring approximately 30 mm in total height. We then placed the cell phone camera approximately 80 mm above the droplet, and captured pictures of the water droplets on a solid white cell phone image.

**Fig. 1.**
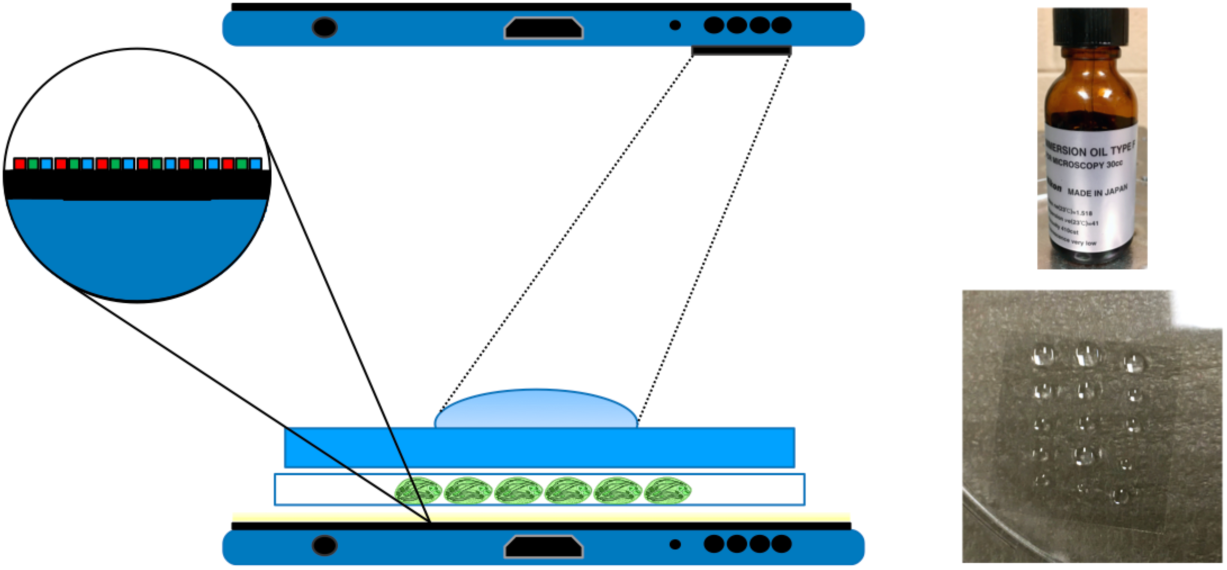
Schematic illustration of a cell phone being used to capture an image of a water droplet that is being used to magnify a biological sample. Use of pixels of known sizes is utilized to quantify the optical magnification of the droplet.

We imported the captured images into ImageJ and measured the size of amplified pixels through each droplet lens. The size of the pixel was defined as the distance between adjacent red, green, or blue pixels near the center of the droplet. The dimension of the image was calibrated by the width of the cell phone screen, which was obtained from the manufacturer’s specifications. The physical size of each pixel was also derived from the pixels per inch (PPI) data from the manufacturer. We then used our magnified pixel values to calculate the magnification factor that each droplet produced and plotted the magnification factor as a function of the droplet volume. For iPhone Xs, we determined the the pixel dimension to be 55.46 µm and the screen width to be 6.22 cm. For the Huawei Honor 7X, we determined the pixel dimension to be 62.41 µm and the screen width to be 6.70 cm.

### 2.2 USAF resolution target analysis

In order to study the resolution of an immersion oil droplet, we used a white light source and a Huawei Honor 7X cell phone to capture images of an oil droplet magnifying a Positive 1951 USAF test target (Thorlabs R1DS1P).

### 2.3 Making of cooking oil droplets

To account for the simplicity of the experiment, we used cooking oil as a source of magnification. We obtained consumer-grade corn oil, canola oil, and corn oil.

### 2.4 Imaging biological samples using immersion oil

After we measured the resolution of the oil droplets, we used a white light source to illuminate two biological slides containing an onion epidermis and a zea stem cross section. We used an oil droplet to magnify the biological samples and captured images of the magnified biological samples using the Huawei Honor 7X cell phone. The cell phone images were compared with images taken using a Plan Apo λ 20x NA 0.75 objective on a Nikon Eclipse Ti-E2 microscope and a Photometric Prime 95B back-illuminated sCMOS camera.

### 2.5 Comparison of smartphone images obtained using immersion and cooking oils

Once we compared the images we obtained using immersion oil to those captured with the Nikon microscope, we prepared a series of 1-5 µL droplets of immersion, canola, olive, and corn oil on glass coverslips. We then captured images of the zea stem cross section and onion epidermis using each set of oil droplets.

## 3. Results

### 3.1 Oil droplets are more resistant to evaporation

Using a cellphone camera to acquire magnified images through a plano-convex lens formed by a water droplet practically constitutes a two-lens system. When using water as a droplet lens, evaporation causes constant change to the radius of curvature and effective focal length of the droplet. We compared the evaporation rates of droplets made of water, immersion oil, and corn oil on a glass coverslip. After 20 minutes at room temperature in an indoor laboratory setting, water droplets smaller than 5 µl completely evaporated, whereas both immersion and corn oil droplets maintained their shape and volume (Fig 2).

**Fig. 2.**
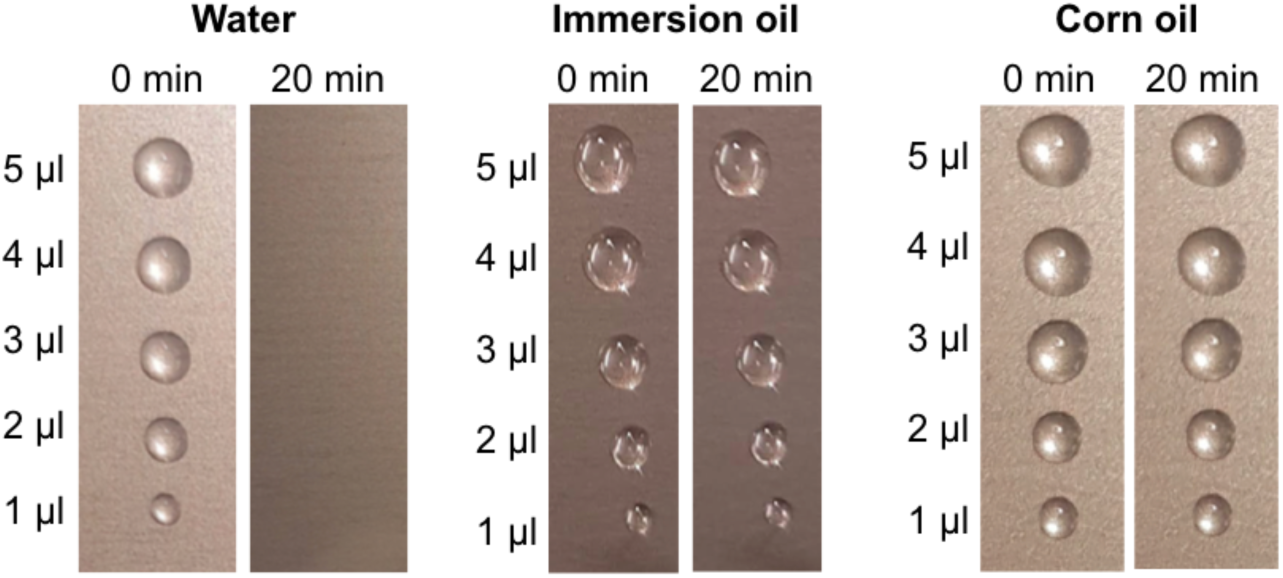
Comparison of droplets made of water, immersion oil, and corn oil at room temperature for 20 minutes.

### 3.2 Characterizing optical resolution using screen pixels and a resolution target

We utilized known sizes of cell phone screen pixels to quantify the optical magnification. Fig 3A illustrates the amplified images of pixels on an iPhone Xs screen through the 5 µl and 2 µl droplets. Optical amplification of water and oil droplets on the coverglass was found to be similar in the range between 3 and 5x when the volume of the droplet is greater than 3 µl (Fig 3B). Smaller oil droplets exhibited much higher magnification than water droplets. The 2 µl oil droplet achieved a magnification of 9.4, about 77% higher than the equivalent water droplet. The higher magnification is likely attributed to the higher surface tension of oil. The angle of contact between the oil droplet and glass is much higher than that between the water droplet and glass. This property creates a higher-power positive lens, and along with the lower tendency to evaporate, rendering immersion oil droplets excellent optical elements for amplifying biological specimen. At 1 µl, the spherical aberration is significant from the oil droplet, rendering it challenging to quantify the magnification.

**Fig. 3.**
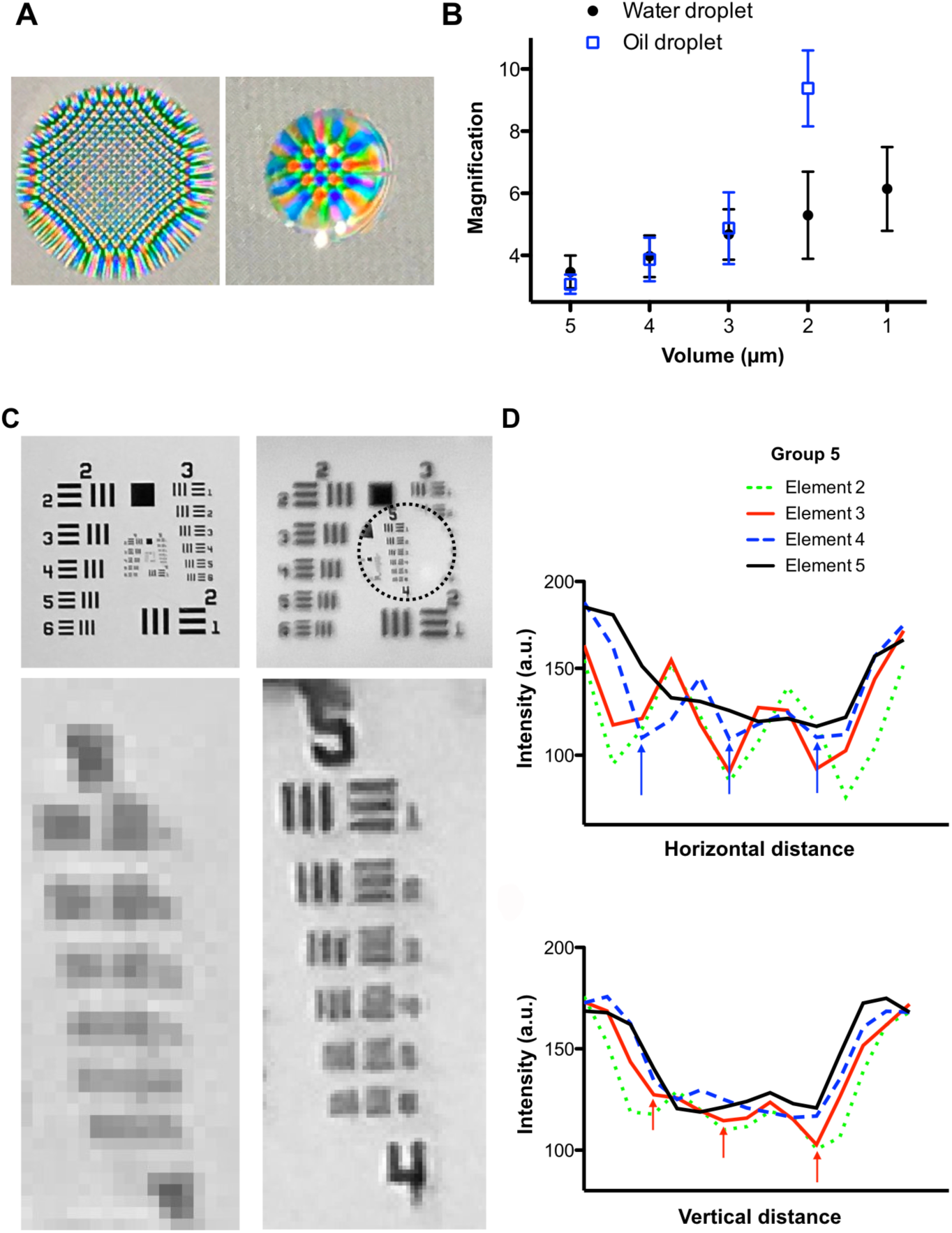
Optical characterization of water and oil droplets. (A-B) Characterizaiton using a cellphone screen. (C-D) Characterizaiton using a USAF resolution target.

To demonstrate the enhancement of optical resolution from the oil droplet, we compared two images of the USAF resolution target with and without the oil droplet. Fig 3C demonstrates a typical image of the resolution target. The right panel demonstrates an image of the same resolution target with an oil droplet outlined by the dotted circle above group 5 elements. While the standard cell phone image failed to resolve any elements within this group, the image magnified by the oil droplet resolved several elements. The line profile plots of the vertical and horizontal elements (Fig 3D) demonstrate the ability of the oil droplet to resolve vertical element 4 and horizontal element 3. The line width of element 3 indicates an optical resolution of 12.40 µm. We note that this resolution is likely to be sensitive to the distance of the object from the droplet as well as the distance between the cell phone camera and the droplet. We demonstrate here a simple and versatile method here to achieve enhanced optical resolution for bioimaging.

### 3.3 Resolving cellular structures using an immersion oil droplet

After we examined the resolution of the oil droplets, we used the oil droplets to obtain images of biological samples, including a zea stem and an onion epidermis. As shown in Fig 4, the images obtained using the iPhone and oil droplets showed the same structures that were obtained using a 20x/0.75 objective on a Nikon Eclipse Ti-E2 inverted microscope. While the Nikon image had a visibly higher resolution than our oil droplet images, the oil droplets enabled us to view biological structures that would have been impossible for us to view using the naked eye. For instance, when the onion epidermis was magnified with the oil droplet, the shapes of individual cells were visible. In the zea stem cross section, the xylem and phloem of the plant vascular structure were also visible, although they were less resolved in the oil droplet image than in the Nikon image.

**Fig. 4.**
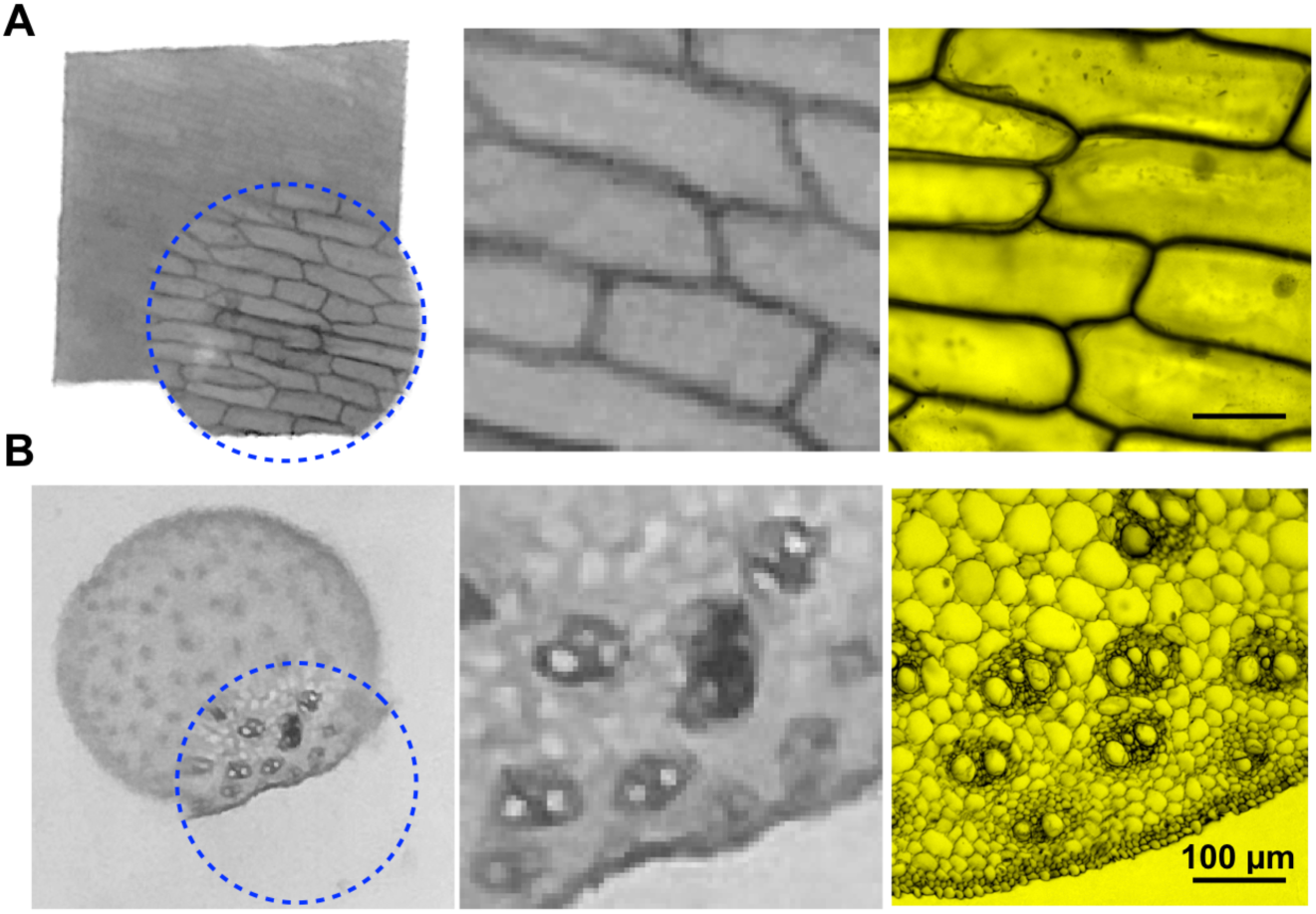
A comparison between magnified oil droplet images and images obtained using a Nikon Eclise Ti-E2 inverted microscope with brightfield illumination and a 20x/0.75 objective. The cell phone images were taken by illuminating an onion epidermis (top) and zea stem cross-section (bottom) with a white light source and using an oil droplet (outlined with a blue circle) to magnify the images. The middle images are high-magnification view of the images on the left, and the images on the right were captured using the Nikon microscope. The biological samples were obtained from AmScope.

### 3.4 Comparing immersion oil and cooking oil droplets for optical imaging

In spite of its ideal properties, the cost of immersion oil may present significant barriers in low-resource settings due to its cost and low accessibility. For this reason, we explored the use of consumer-grade oils, which have historically been used to conduct immersion oil microscopy before synthetic immersion oils were commercially available. Before synthetic oils became the standard for immersion oil microscopy, natural oils such as cedar tree oil and castor oil were typically used ^18^. More recently, castor oil has been used to obtain immersion objective images of lymphocytes in metaphase to produce images comparable to those taken using synthetic immersion oil.

We obtained canola oil, olive oil blend, corn oil, and Nikon immersion oil to evaluate their performance for smartphone microscopy. Two sets of coverslips were prepared for each oil; one coverslip contained unknown volumes of oil and the other coverslip contained known volumes of oil. Approximately 1 mL of each oil was placed into four conical cylindrical centrifuge tubes to be used for preparation of oil droplets. A pipette tip was then used to transfer small amounts of oil from the centrifuge to the coverslips and form three rows of oil droplets with unspecified volumes. After the droplets of unknown volumes were prepared, the centrifuge tubes containing each oil were heated to 37°C for ten minutes using a hot water bath to reduce the viscosity of the oils. We then prepared one coverslip for each heated oil onto which we placed droplets ranging from 2-4 µL using a micropipette.

Once our oil droplets were prepared, we used them to obtain a series of images of the zea stem and onion epidermis samples. Roughly, similar color and resolution were achieved using cooking oil droplets compared to immersion oil droplets. Generally, images that were taken using 4 µL droplets were less magnified and less resolved than images taken using 3 µL and 2 µL droplets, as expected (Fig 5). However, there were several images that deviated from this trend; for instance, the 3 µL and 4 µL immersion oil droplets appeared to provide approximately equal resolution.

**Fig. 5.**
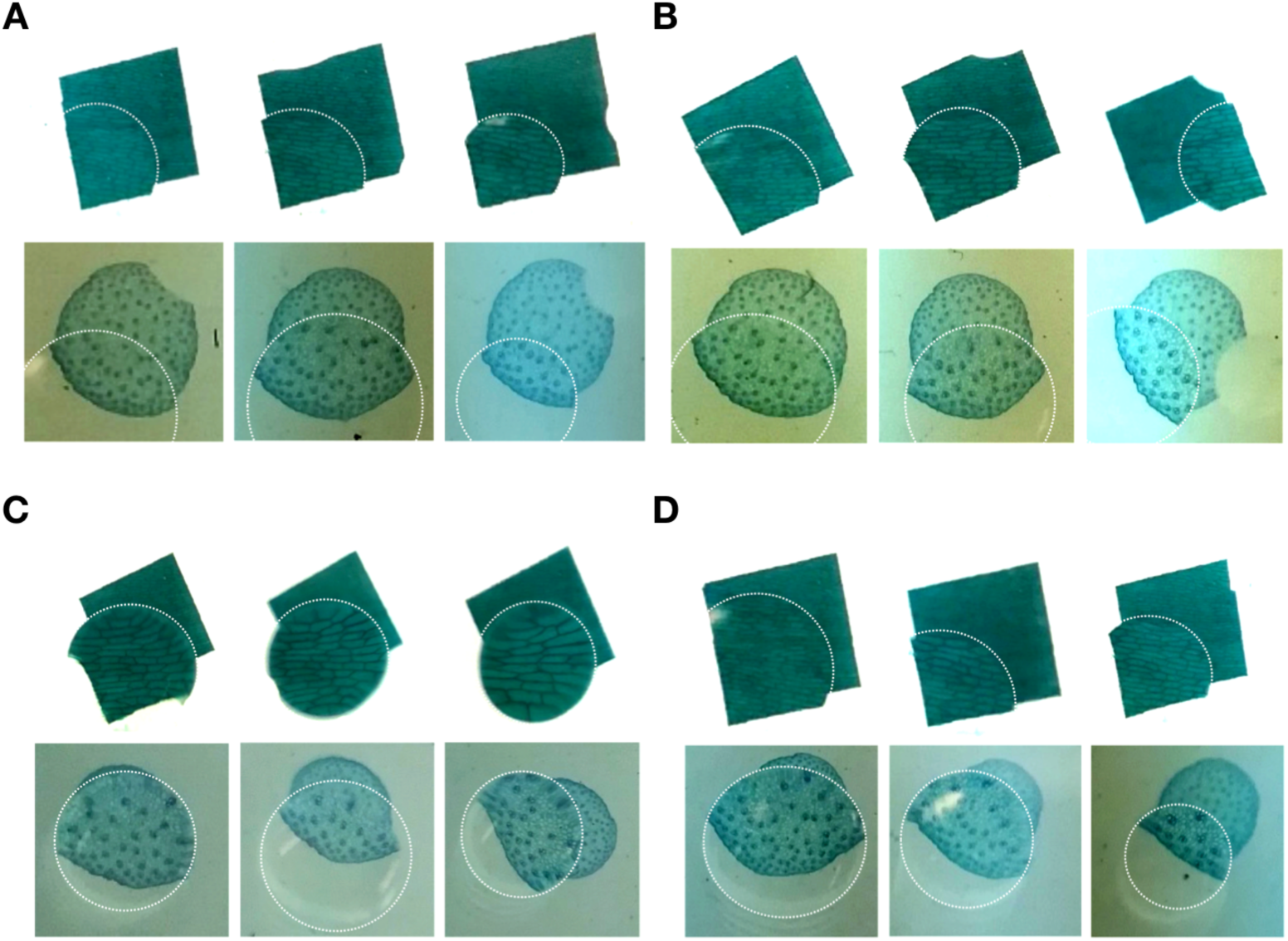
Imaging using cooking oil droplets of 4, 3, and 2 µL in volume, from left to right. A) Immersion oil, B) corn oil, C) canola oil, D) olive oil.

## Discussion

In the future, liquid droplets could be used to design flexible, low-cost microscopy systems for educational or diagnostic purposes. Immersion oil, in particular, could be used as a somewhat more permanent lens to view microscopic structures, since it does not evaporate nearly as quickly and has the same refractive index as glass. As can be seen in Fig 2, while the row of water droplets evaporated almost entirely in about twenty minutes, the oil droplets did not decrease in size over the measured time. Therefore, if an oil droplet lens of a specific size was needed for multiple imaging sessions, the same oil droplet lens could be reused during multiple imaging sessions, which would save time and retain the previously used magnification factor. While glass lenses can be expensive and require extensive manufacturing processes, oil droplet lenses only require the use of a coverslip and plastic tool to transfer oil. While we used a micropipette to create our oil droplets, a micropipette is only needed when exact volumes of oil are desired, which is not required to obtain qualitative images of biological samples. Furthermore, since several oil droplets can be made quickly, it is possible to easily prepare a series of droplets that increase in magnification with either a wide or narrow magnification range, depending on what type of images are being captured. Therefore, microscopy using oil droplets and cell phones for sample illumination and image capture are promising yet simple tools that could be refined in the future to potentially conduct quantitative and qualitative analyses of biological samples.

In our experiments, we demonstrated that cooking oil may be a useful, cost-effective method of obtaining images of biological samples in a low-resource environment. We initially showed that images obtained using immersion oil droplets are resolved up to 12.4 microns, which is a significant improvement over the naked eye. We then compared the immersion oil droplet images to the images we obtained using a conventional Nikon microscope, which showed that the structures we viewed using immersion oil lenses were consistent with those seen using conventional lenses. In order to address the issue of cost-effectiveness in a low-resource setting, we experimented with the use of cooking oils as oil droplet lenses, since cooking oils such as castor oil have a history of being used in immersion oil because their refractive indices are close to that of glass and of synthetic immersion oil. We found that images obtained using cooking oil droplets were similar to those obtained using immersion oil droplets, and that oil droplets could be prepared with or without the use of a micropipette. While we used a pipette tip to transfer the oils when preparing droplets of non-specified volumes, a less expensive alternative could be used in a low-resource setting, such as a plastic stick. Although we were able to produce useful images through the use of cooking oil droplets, the resolution and focus levels that we achieved were, at times, inconsistent. This was likely due to settings that are pre-programmed into cell phones that make them less ideal imaging tools for scientific applications. This obstacle is a potential avenue for improvement that may be explored in future studies.

While cell phones are designed to be cost-effective and simple to use, cell phone imaging often sacrifices quality and versatility for simplicity. Studies have shown that cell phones have certain built-in limitations that reduce their accuracy and hinder their use as a quantitative tool for diagnostic microscopy ^19^. One of the key differences between scientific and cell phone microscopes is that scientific imaging systems allow users to have full control over the camera or microscope settings, whereas cell phone microscopes are programmed with default settings, some of which cannot be changed easily. Many smartphones use auto-focus, which can lead to inaccurate size quantification in microscopy because it can impact magnification by changing the apparent size of a structure as much as 6% ^19^. Another feature that makes it more challenging to conduct cell phone microscopy is the automatic image processing that is programmed into cell phones, such as noise reduction and image compression, which are useful in generating desirable images in cell phone photography, but often lead to a loss of information and inconsistent imaging, which can lead to inaccurate quantitative analysis in cell phone microscopy ^19^. For instance, the image quality in Figs 3-5 is subject to the image processing algorithm within the cell phone. We expect the use of calibration standard, such as the known pixel sizes of a cell phone screen, to mitigate the uncertainties and enable quantitative imaging.

There are several challenges that may emerge when working with cooking oil droplets; for instance in an outdoor environment, droplets may attract dust particles, which could potentially deteriorate image quality. While our method is only aimed to be utilized in the short-term, ranging from a few minutes to several days, it may be useful to take measures to account for possible contamination from dust or other particulates. This may be achieved by enclosing coverslips with oil droplets in a container, such as a petri dish, to prevent contamination. Another potential issue is that some vegetable oils such as sesame oil and olive oil are naturally colored, which may impact apparent color of the biological samples in the images obtained using a cell phone. There was no noticeable difference in color between the images we obtained using olive oil and other types of vegetable oil. While we did not see a major color difference due to natural oil colors, it may be valuable to conduct smartphone imaging experiments comparing different brands of several types of vegetable oils to see if there is a color difference between them.

In the future, it may also be useful to consider cost-efficient methods that would increase the tunability of vegetable oil droplet lenses, which would provide more flexibility in low-resource cell phone microscopy. For instance, surface properties such as polarity, can affect the wetting and shape of the droplet. These surface properties can be exploited in future studies to explore the tunability of cooking oil droplet lenses. Taken together, the ease of operation in droplet-based bioimaging will extend the discoveries from medical and biology researcher to the hands of field workers and educators.

## Conclusion

We present an accessible imaging method using evaporation-resistant oil droplets and the cellphone camera. The attenable optical resolution enables direct observation of cellular structures in plant tissue samples. We further demonstrate the applicability of using household oils for optical imaging. Combined with the versatility of capturing and sending digital images through the mobile network, our study lays the groundwork for an attractive optical technology for improving healthcare in low-resource settings with a minimal footprint.

## References

1. Smith, Z. J. et al. Cell-phone-based platform for biomedical device development and education applications. PLoS One 6, e17150 (2011).

2. Orth, A., Wilson, E. R., Thompson, J. G. & Gibson, B. C. A dual-mode mobile phone microscope using the onboard camera flash and ambient light. Sci. Rep. 8, 3298 (2018).

3. Dai, B. et al. Colour compound lenses for a portable fluorescence microscope. Light: Science & Applications vol. 8 (2019).

4. Pirnstill, C. W. & Coté, G. L. Malaria Diagnosis Using a Mobile Phone Polarized Microscope. Sci. Rep. 5, 13368 (2015).

5. Jung, D. et al. Smartphone-based multi-contrast microscope using color-multiplexed illumination. Sci. Rep. 7, 7564 (2017).

6. Kim, J.-H., Joo, H.-G., Kim, T.-H. & Ju, Y.-G. A smartphone-based fluorescence microscope utilizing an external phone camera lens module. BioChip Journal vol. 9 285–292 (2015).

7. Zhu, H., Yaglidere, O., Su, T.-W., Tseng, D. & Ozcan, A. Wide-field fluorescent microscopy on a cell-phone. Conf. Proc. IEEE Eng. Med. Biol. Soc. 2011, 6801–6804 (2011).

8. Bellina, L. & Missoni, E. Mobile cell-phones (M-phones) in telemicroscopy: increasing connectivity of isolated laboratories. Diagn. Pathol. 4, 19 (2009).

9. Agbana, T. E. et al. Imaging & identification of malaria parasites using cellphone microscope with a ball lens. PLOS ONE vol. 13 e0205020 (2018).

10. Kobori, Y., Pfanner, P., Prins, G. S. & Niederberger, C. Novel device for male infertility screening with single-ball lens microscope and smartphone. Fertility and Sterility vol. 106 574–578 (2016).

11. Sung, Y.-L., Jeang, J., Lee, C.-H. & Shih, W.-C. Fabricating optical lenses by inkjet printing and heat-assisted in situ curing of polydimethylsiloxane for smartphone microscopy. J. Biomed. Opt. 20, 047005 (2015).

12. Zeng, X. & Jiang, H. Liquid tunable microlenses based on MEMS techniques. Journal of Physics D: Applied Physics vol. 46 323001 (2013).

13. Myint, H. H., Marpaung, A. M., Kurniawan, H., Hattori, H. & Kagawa, K. Water droplet lens microscope and microphotographs. Physics Education vol. 36 97–101 (2001).

14. Chowdhury, F. A. & Chau, K. J. Microscopy using water droplets. MOEMS and Miniaturized Systems XI (2012) doi:10.1117/12.906949.

15. Refractive Index List of Common Household Liquids. International Gem Society https://www.gemsociety.org/article/refractive-index-list-of-common-household-liquids/.

16. Arya, S. S., Ramanujam, S. & Vijayaraghavan, P. K. Refractive index as an objective method for evaluation of rancidity in edible oils and fats. Journal of the American Oil Chemists’ Society vol. 46 28–30 (1969).

17. [No title]. https://www.ijaar.org/articles/Volume4-Number4/Sciences-Technology-Engineering/ijaar-ste-v4n4-apr18-p42.pdf.

18. Reddy, D. J. Substitute for Cedar Wood Oil for Oil Immersion Work. Ind. Med. Gaz. 79, 565 (1944).

19. Skandarajah, A., Reber, C. D., Switz, N. A. & Fletcher, D. A. Quantitative imaging with a mobile phone microscope. PLoS One 9, e96906 (2014).

